# Efficient storage and regression computation for population-scale genome sequencing studies

**DOI:** 10.1101/2024.04.11.589062

**Authors:** Manuel A. Rivas, Christopher Chang

## Abstract

In the era of big data in human genetics, large-scale biobanks aggregating genetic data from diverse populations have emerged as important for advancing our understanding of human health and disease. However, the computational and storage demands of whole genome sequencing (WGS) studies pose significant challenges, especially for researchers from underfunded institutions or developing countries, creating a disparity in research capabilities. We introduce new approaches that significantly enhance computational efficiency and reduce data storage requirements for WGS studies. By developing algorithms for compressed storage of genetic data, focusing particularly on optimizing the representation of rare variants, and designing regression methods tailored for the scale and complexity of WGS data, we significantly lower computational and storage costs. We integrate our approach into PLINK 2.0. The implementation demonstrates considerable reductions in storage space and computational time without compromising analytical accuracy, as evidenced by the application to the AllofUs project data. We optimized the runtime of an exome-wide association analysis involving 19.4 million variants and the body mass index phenotype of 125,077 individuals, reducing it from 695.35 minutes (approximately 11.5 hours) on a single machine to just 1.57 minutes using 30 GB of memory and 50 threads (or 8.67 minutes with 4 threads). Additionally, we extended this approach to support multi-phenotype analyses. We anticipate that our approach will enable researchers across the globe to unlock the potential of population biobanks, accelerating the pace of discoveries that can improve our understanding of human health and disease.

## I. INTRODUCTION

The era of big data in human genetics has ushered new opportunities for understanding human health and disease and generating effective therapeutic hypotheses^1–3^. Large-scale biobanks, aggregating genetic data from diverse populations, have become a cornerstone for this new wave of research^4–8^. These repositories offer new insights into the genetic underpinnings of diseases, facilitating discoveries. However, the shift towards leveraging virtual machine workbenches for genetic analysis across these population biobanks presents a new set of challenges. The computational intensity and voluminous data associated with whole genome sequencing (WGS) studies require significant resources, imposing substantial computational and storage costs. This burden renders advanced genetic analysis inaccessible to a broad spectrum of researchers, particularly those from underfunded institutions or developing countries, thereby creating a disparity in research capabilities and slowing the pace of scientific discovery.

Recognizing these barriers, we introduce new approaches that address the dual challenges of computational efficiency and data storage. We have developed algorithms that facilitate compressed storage of genetic data, significantly reducing the storage requirements for WGS studies. This compressed representation leverages patterns within genetic variants focused on optimizing representation of rare variants, enabling a more compact representation of genetic data without sacrificing data integrity^9^. By diminishing the storage footprint, our approach lowers the cost associated with data storage on cloud platforms and other virtual machine environments, making large-scale genetic data more manageable and accessible.

Furthermore, our work extends beyond data compression to tackle the computational inefficiencies inherent in current genetic analysis methods. Traditional regression techniques, while robust, are not optimized for the scale and complexity of data generated by WGS studies. Our regression approaches are designed specifically to address the unique challenges presented by case-control, quantitative, and multi-phenotype analysis where a large fraction (*>* 99.99%) of the greater than 1 billion of genetic variants discovered from whole genome sequencing studies are rare (i.e. minor allele frequency less than 1% in the population)^10^. These methods utilize sparse computation techniques to enhance processing speed without compromising analytical accuracy. By focusing computational resources only on the most relevant genetic data, i.e. individuals that carry the rare variant, we can expedite the analysis process, enabling researchers to conduct more complex and comprehensive studies with limited resources.

Our implementation of these algorithms in PLINK 2.0 integrates seamlessly with existing virtual machine workbenches, offering an accessible and user-friendly tool for genetic researchers.

### II. METHODS

Significant optimizations have been introduced to the computational processes involved in Generalized Linear Model (GLM) analysis to enhance efficiency, accuracy, and memory management^11–13^ addressing the computational and storage challenges.

PGEN is PLINK 2’s binary genotype file format. It introduces a sparse representation for rare variants: in the PLINK 1 binary format, a variant with a single alternate-allele observation out of 400000 samples would require the same 100000 bytes as any other variant in the file, but the PGEN representation typically requires only 4 bytes in the file header and 5 bytes in the body (logically representing “all genotype values are 0 except sample *x*, which has genotype value *y*”). Importantly, it is possible to perform computation directly on this sparse representation; in contrast, BGEN and BCF compress rare variants but force readers to fully decompress them. PGEN also includes a simple form of LD-based compression, and provides ways to directly represent multiallelic variants, phase, and dosage information, which are outside the scope of this document; see https://github.com/chrchang/plink-ng/tree/master/pgen_spec for full details.

PLINK 2.0’s `--glm` command executes per-variant linear-or logistic/Firth-regressions between a phenotype vector *y* and a predictor matrix **X**, where **X** includes genotype and intercept columns, and usually includes some covariates (e.g. top principal components). We discuss sparse-genotype optimizations implemented in the quantitative phenotype linear regression code here, some of which are also applicable to binary phenotypes.

Denote the number of samples by *n*, and the number of predictors by *p*, so *y* is a *n ×* 1 vector and **X** is a *n × p* matrix. The least squares linear regression solution is 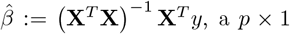 vector. The residual sum of squares can be expressed as 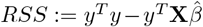. The p-values and standard errors reported by `--glm` on a quantitative phenotype are simple functions of 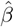 and *RSS*. Ordinarily, `--glm` performs this computation by filling floating-point arrays with the contents of *y* and **X**, and using BLAS/LAPACK^15^ routines to compute 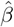 and *RSS*. When multiple phenotypes covering the exact same samples are provided, PLINK 2.0 processes them simultaneously (*y* is replaced with a wider **Y** matrix) if there is sufficient memory.

The first implemented sparse-genotype optimization addressed the no-missing-genotype, no-dosage case.

Write **X** as the block matrix (**X**_*c*_ *x*_*g*_), where **X**_*c*_ contains all non-genotype predictors and *x*_*g*_ is the genotype column vector. Then, 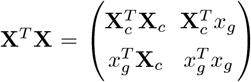. The inverse of a 2×2 block matrix 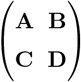 is, when **A** and **F** := **D***−***CA**^*−*1^**B** are invertible, 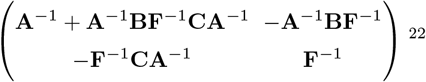. We compute 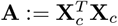 and **A**^*−*1^ before any variants are processed. Then, for a sparse variant with *k* nonzero values in *x*_*g*_, we can finish evaluating (**X**^*T*^ **X**)^*−*1^ by:

1. Evaluating 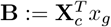 (and **C** := **B**^*T*^). This requires *𝒪*(*kp*) arithmetic operations.
2. Evaluating 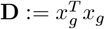. This requires *𝒪*(*k*) arithmetic operations.
3. Evaluating **F** := **D** *−* **CA**^*−*1^**B**. This requires *𝒪*(*p*^2^) arithmetic operations.
4. Since **F** is 1×1, **F**^*−*1^ can then be evaluated by taking a single reciprocal.
5. The remaining matrix-addition and vector-matrix-multiply steps require *𝒪*(*p*^2^) arithmetic operations.

This adds up to *𝒪*(*p*(*k* + *p*)) arithmetic operations. 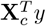 can be computed in advance, so evaluation of **X**^*T*^ *y* requires only *𝒪*(*k* + *p*) arithmetic operations. Finally, *y*^*T*^ *y* can also be computed in advance, so the cost of determining 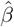 and *RSS* is potentially still *𝒪*(*p*(*k* + *p*)) arithmetic operations. It is unnecessary to pay the cost of populating a full floating-point **X** matrix, or the higher cost of LAPACK operations on it.

Missing genotypes were not supported by this code path, since they raise the cost to *𝒪*(*p*^2^(*k* + *p*)): the precomputed 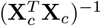 value cannot be used. To keep the code simpler, dosages also weren’t supported; it was expected that dosage-containing genotype columns were not likely to be very sparse.

We note that refitting the covariates separately for every single variant is unnecessary, especially if we are to apply the regression across billions of genetic variants. This approach has been used historically in genome sequencing analysis of quantitative traits as described in the GoT2D project^16^. We introduce the `qt-residualize` modifier in `PLINK 2.0`, which performs a single regression on the covariates upfront, and then passes the phenotype residuals and no covariates to the main per-variant-linear regression loop described above.

When `qt-residualize` is in effect, *p* is reduced to 2 (genotype and intercept), so missing genotypes are no longer much more expensive for a sparse algorithm to handle than ordinary nonzero genotype values. So we have implemented a second sparse-genotype code path covering the *p*=2, no-dosage case: it subtracts from 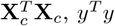 and 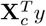 every time a missing genotype is encountered, and otherwise follows the same mathematical approach as the first sparse algorithm. This code path takes full advantage of the PGEN format: when a rare variant is represented as a sparse list of nonzero (sample ID, genotype value) tuples, it directly processes such a list instead of forcing it to be expanded to an array of *n* 2-bit values.

## III. RESULTS

Given that the virtual research workbench (where the AllofUs data resides) makes available a limited set of storage (Gb) and computational resources (RAM, CPUs) we asked whether we could compress the sequencing data substantially beyond that provided in the v7 release. We started with the exome sequencing data which has a mixture of common and rare variants as proof of principle of our approach. We anticipate that the approach we take here will generalize to whole genome sequencing data. We compare compression of genetic variant genotype data from the exome sequencing subset of the AllofUs data. We find that 1) the initial PLINK BED/FAM/BIM binary file format requires approximately 2Tb of storage; 2) Hail SplitMT and MultiMT requires 409Gb and 208Gb, respectively; 3) VCF file requires 403Gb; 4) BGEN file requires 165Gb; and 5) PGEN file requires 39.0Gb. PGEN is 98% compressed compared to BED files, 90% compared to VCF files, and 77% compressed compared to BGEN (58.5% if starting from PGEN to BGEN conversion) (see Figure 1).

**FIG 1:**
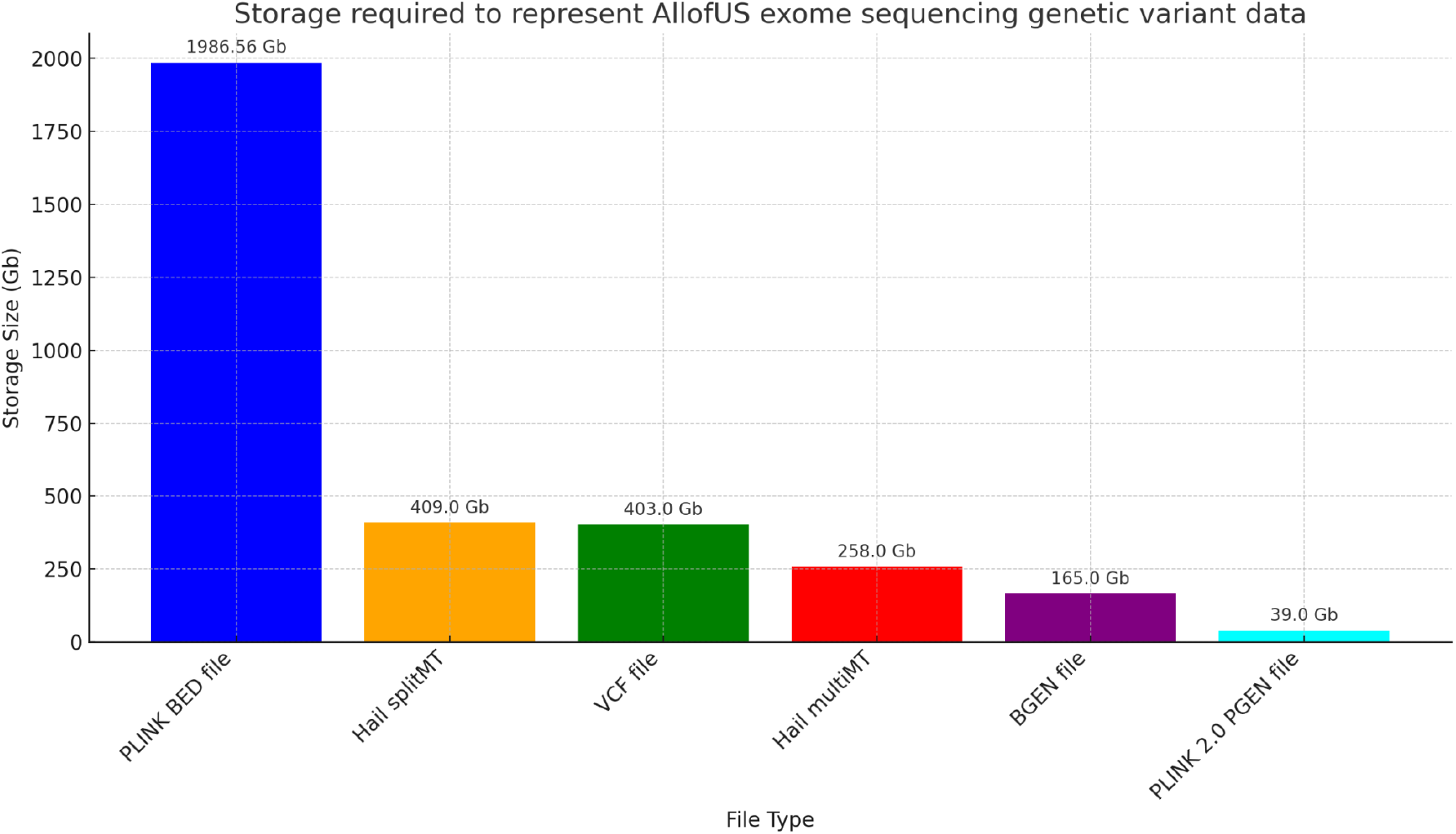
Comparison of storage allocated to exome sequencing data in AllofUs project. Storage required to represent AllofUs exome sequencing genetic variant data for a range of file types. Each bar denotes the storage size, with values provided on top for clarity. This includes PLINK binary BED file, Hail splitMT and Hail multiMT files, VCF file, BGEN, and PLINK 2.0 PGEN file with sparse variant representation.

To compare computational efficiency we ran univariate GLM regression across 19.33 million genetic variants from the exome sequencing data in 125,077 individuals with body mass index (BMI) phenotype. We find that our standard approach for GLM using PLINK PGEN files implemented in PLINK 2.0 requires approximately 695 minutes using 50 threads, which is approximately 12 hours. We applied an early draft of the qt-residualize mode described in the methods section, and found that it reduces the time to complete the regression analysis to 10 minutes using 50 threads. We added the PGEN sparse-list optimization described at the end of the methods section and found that this additional step reduces the time to complete the regression analysis to 8.67 minutes using only 4 threads and 1.57 minutes if we use 50 threads (see Figure 2).

**FIG 2:**
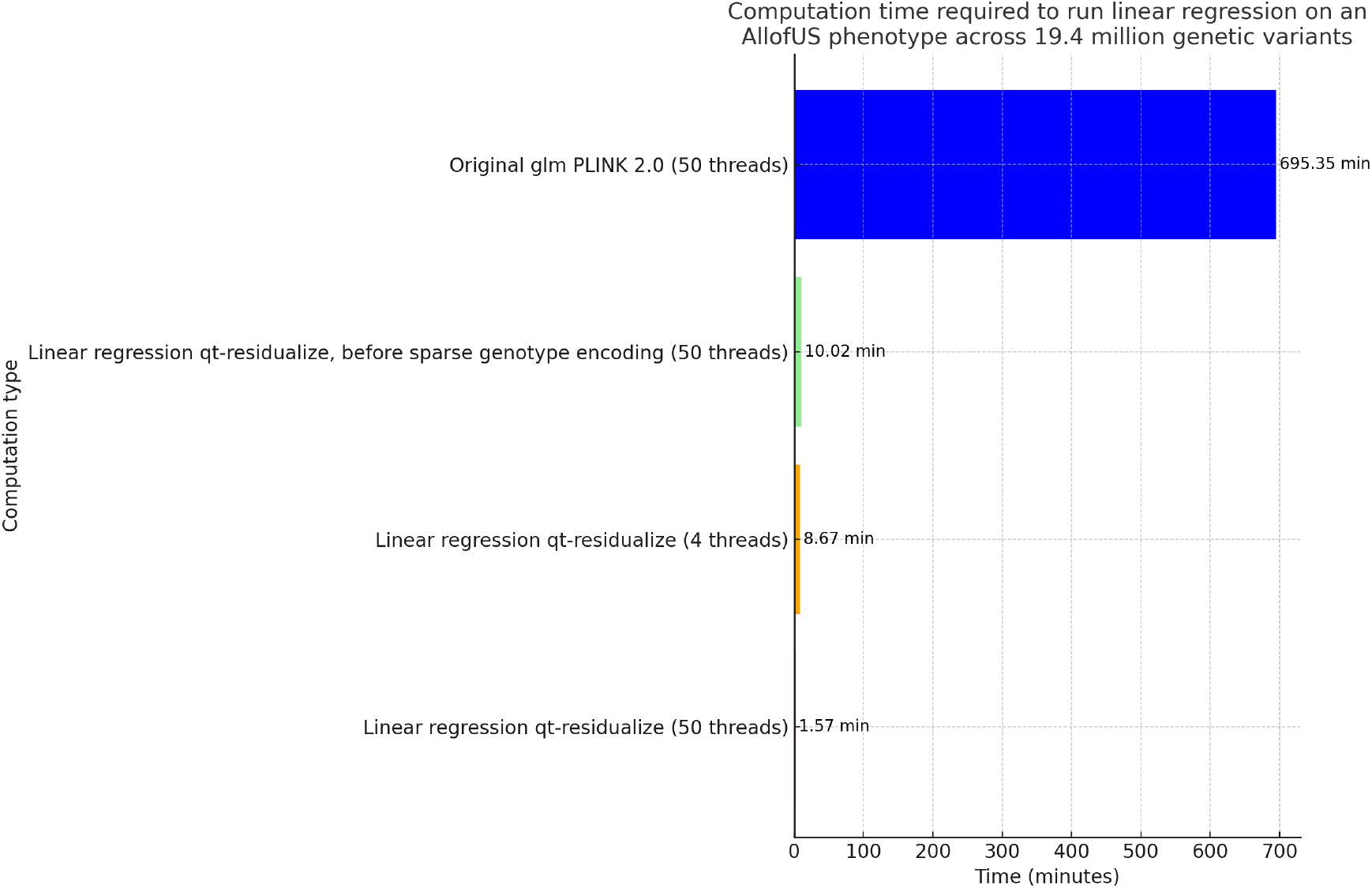
Comparison of regression computational time. Computation time required to run regression on AllofUs body mass index (BMI) phenotype using 18 covariates including age, sex, and 16 principal components of the genetic data, across 19.4 million genetic variants in 125,077 individuals, for different computation scenarios. Each bar displays the time in minutes, with values provided on top for immediate reference. We highlight the significant efficiency gains achieved through optimizing computation methods and sparse genotype data representation for rare variants with PLINK 2.0.

We wondered whether the `qt-residualize` mode introduces variation in P-values we obtain in our regressions across the approximately 20 million variants. Overall, we find that the -log10(P-values) using the original approach, which required a runtime of approximately 12 hours with 50 threads, and the -log10(P-value) using the optimized approach, which required a runtime of approximately 1.57 minutes has an R^2^ of 0.999986 and slope of 0.99937 (see Figure 3).

**FIG 3:**
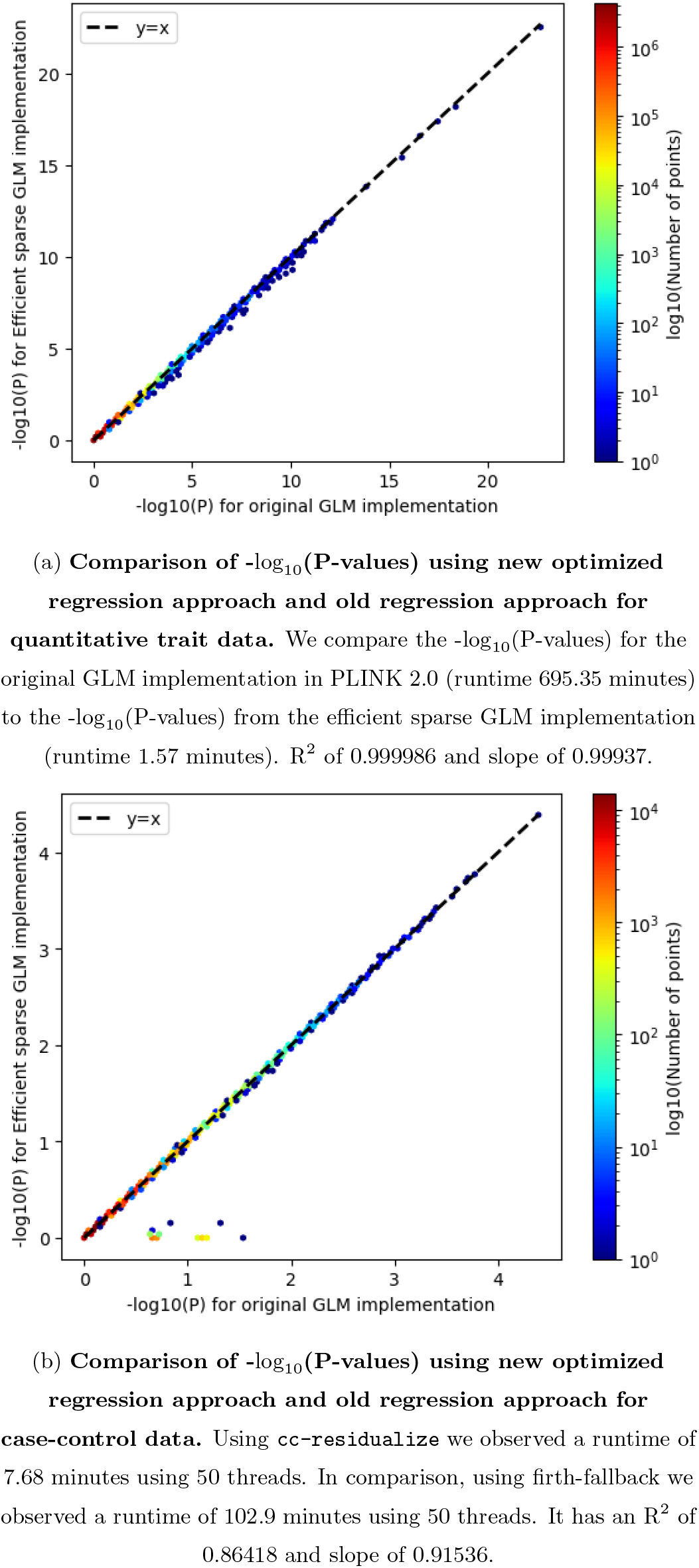
P-value comparison for quantitative and binary data.

We also introduced the `cc-residualize` mode for binary case-control data. We compared computational efficiency gains to firth-fallback implementation in PLINK2. When we analyzed the type 2 diabetes phenotype (N = 16,481 cases and 108,596 controls) across 349,069 variants in chromosome 21 from the AllofUS dataset we observed computational gains. Using `cc-residualize` we observed a runtime of 7.68 minutes using 50 threads. In comparison, using firth-fallback we observed a runtime of 102.9 minutes using 50 threads. It has an R^2^ of 0.86418 and slope of 0.91536 (see Figure 3).

Finally, we asked whether we could apply the same approach to multi-phenotype analysis as then we only need a computation across a **Y** matrix of size *N × k* for the *N* rare variant carriers and the *k* phenotypes corresponding to those individuals. We started with 50 phenotypes using a single virtual machine with 30 Gb of memory (RAM) and 50 threads. The genome-wide association analysis across 19.4 million genetic variants took 52 minutes and 38 seconds. Given that a large number of measurements (assuming the number of phenotypes *>* 1000) obtained from deeply phenotyped population biobanks are correlated, it is natural to ask whether you can start by taking the singular value decomposition (SVD) of **Y** to the minimal number of components that explain a large fraction of the variance in **Y**. However, we acknowledge that results on phenotype components or their linear combinations can pose challenges for interpretation. Despite this, projecting genetic associations onto trait-specific and shared components can offer insights into both unique and overlapping genetic architectures, as highlighted in prior studies where we assessed the components of genetic associations from UK Biobank across 2,138 phenotypes ^17,18^. In our implementation we see that computing regressions across 50 phenotypes (treated to be the same phenotype) takes 50 times as long as when analyzing it as a single phenotype. Given that it is the same phenotype replicated 50 times the matrix **Y** can easily be summarized by a single component. We introduce the `--pheno-svd` flag to preprocess complete rows of phenotype matrix **Y** with variance explained as a parameter to get a lower rank matrix prior to running association analysis (along with a worked missing-phenotype-imputation example in the documentation), and find that for our example we recover the initial 2 minutes required to compute across the 50 phenotypes (a special case where this decomposition aids computation tremendously)^19^.

## IV. DISCUSSION

Our approach, designed to alleviate the computational and storage burdens associated with large-scale genome sequencing studies, sets a new precedent for efficiency and accessibility in genetic research.

Reprocessing and reanalyzing genetic data from biobanks become significantly more feasible. Given the dynamic nature of genetic research, where interpretations and methodologies continually evolve, the ability to efficiently revisit and analyze existing datasets is invaluable.

Researchers can apply new hypotheses to previously unmanageable datasets, uncovering insights that were previously obscured by computational limitations. This not only maximizes the utility of existing genetic data but also accelerates the pace of discovery by facilitating iterative research processes.

Furthermore, our approach will expedite the visualization of genetic inferences, a critical aspect of genetic research that aids in the interpretation and communication of complex data^20^. The ability to succinctly visualize relationships within genetic data as portrayed in Global Biobank Engine (GBE), gnomAD browsers, and other phenotype web browsers can illuminate patterns and connections that are crucial for understanding genetic influences on health and disease^21^. Efficient data handling and analysis techniques can generate more insightful visualizations, enabling researchers to navigate and interpret genetic landscapes with greater clarity and confidence.

As the field of genetic research progresses, methods that enhance computational efficiency and reduce storage needs will become increasingly important. These approaches will be key to fully leverage the insights from genome sequencing studies, thereby leading to a more productive and innovative research environment.

## V. ACKNOWLEDGEMENTS

M.A.R. is in part supported by National Human Genome Research Institute (NHGRI) under award R01HG010140, and by the National Institutes of Mental Health (NIMH) under award R01MH124244 both of the National Institutes of Health (NIH).

Some of the computing used was done in the AllofUs virtual workbench. The content is solely the responsibility of the authors and does not necessarily represent the official views of the funding agencies; funders had no role in study design, data collection and analysis, decision to publish, or preparation of the manuscript.

This research has been conducted using the AllofUs resource using application, “Effective therapeutic hypotheses” (PI: Manuel A. Rivas).

